## Appendix for "Automatically tracking feeding behavior in populations of foraging *C. elegans*"

**1. Collision detection and effective number of independent animals in the field-of-view**

PharaGlow detects all fluorescent objects in the field of view. The experimenter should choose a minimal and maximal size to allow only tracking objects that are single worms. If animals touch, an object that is larger than the maximally allowed size (maxSize) will be detected. In this case, the program automatically re-segments the region and attempts to separate the large object into multiple smaller objects using repeated filtering and thresholding. If this process is successful and two or more objects of the correct size can be separated, the program will continue tracking these animals. This approach is successful when animals touch, but do not overlap. We are unable to resolve crossings where animals are physically overlapping.

To determine the effect of collisions on tracking, we determine the number of detected objects in the field-of-view and compare it to the number of tracks obtained. Analysis of the track duration shows that average tracks are two min, and this duration depends on the velocity (Fig. S3.6A). We find that these two measures correlate well, supporting the view that animals are not frequently broken into small tracklets (Fig. S3.6B).

**2. Computational cost and scaling**

We benchmarked the performance and scaling of the software using perfplot. Table S1 details the pure computation time as run on a 1000 frame demo recording (available for download at the data repository). We find that parallelization improves performance for the object detection for up to 8 workers, and continues improving for >16 workers in the segmentation step. In our implementation, this option is already provided based on the python package multiprocessing. As the different steps depend on the details of the imaging, we have decided to report processing time per step.

1. Object detection

   The object detection step uses a full frame and does masking and object detection for each individual object. For this case, computational cost scales with the number of frames and can be easily parallelized to enable a faster speed. The average single-core computation time per frame is 300 ms, which includes I/O, as we employ lazy data loading, which allows analyses of data that are much larger than the RAM available. Of note, this step can be omitted if our acquisition software is used (see Methods) as single worms are segmented already during acquisition.

2. Tracking and trajectory interpolation
   The tracking step is based on trackpy and here the scaling depends on the search range (how far can an object move between frames) and the memory (how many subsequent frames can an object be unobserved). Typically, this step is much faster than the other two as it does not handle large I/O or image processing.

3. Segmentation, centerline detection, straightening, and pump detection
   In this step, the previously detected images of detected pharynx are further processed. The total compute time here depends on the product between the number of objects and the number of recording frames. We therefore provide a per-object assessment of the processing time.

**Table S1: Benchmarking of the typical computing times.** Full frames are the multi-animal images with 3088x2064 px. Mini-frames are small regions of interest with one animal (Fig. 1D).

| | **Compute time*** *Intel(R) Xeon(R) Gold 6230 CPU @ 2.10GHz | | |
|---|---|---|---|
| **Step** | 1 worker | 4 workers | 16 workers |
| Object detection | 186.6 ms / full frame | 81.2 ms / full frame | 57.4 ms / full frame |
| Tracking and trajectory interpolation | 1 ms / miniframe | 1 ms / miniframe | 1 ms / miniframe |
| Segmentation etc. | 378 ms / mini-frame | 104 ms / miniframe | 32 ms / miniframe |
| **Typical recording** 150 worm minutes of data (9000 full frames, 30 worms) | **1.2 days** | **8.2 h** | **3 h** |

### 3. Effects of light exposure

*C. elegans* are known to sense and react to light by initiating reversals and suppressing pumping. These reactions occur more frequently at short wavelengths and high power densities [1,2,3]. To determine if our imaging conditions affected behavior, we measured the light intensity and the leaving rates of animals during imaging. We used excitation light centered at 500 nm, and measured an effective intensity of only 0.24 mW/mm$^2$ in the field-of-view, 54 times lower than the reported intensity that induces pumping inhibition or spitting[3]. We observed 5-25% of animals leaving the field-of-view during recordings, indicating a mild avoidance reaction which depends on the developmental stage (Fig. S2.3).

To control for photo-toxic effects, we split our developmental cohort into two groups. One group of animals was imaged consecutively at each larval stage (multiple exposures), the other group was left to grow under the same conditions, but only ever imaged once (single exposure). We find that during all larval stages, the behavioral results of the two groups are similar, but not in young adults (Fig. S2.4). For the young adult cohort, the animals that were repeatedly imaged show a higher velocity compared to the single-exposure group, as well as differences in all other behavioral metrics we report. We speculate that this could be due to differences in drying of the plates during repeated imaging, or a possible light-induced chronic effect.

Over longer time scales, exposure to light can reduce the worms' lifespan[4]. To test for chronic photo-toxic effects, we tested the viability of worms after long exposure to 500 nm light. We continuously illuminated 30 young adult GRU101 worms for five hours using the same illumination intensity as our PharaGlow assay (0.24 mW/mm$^2$ at the focal plane). We employed a copper frame to prevent the worms from escaping the illuminated area. A plate not exposed to the 500 nm light was placed on the same bench close to the microscope as our negative control. After five hours, the copper frame was removed and worms were scored for viability both immediately and after overnight recovery in a 20°C incubator. All worms were viable and able to move upon gentle tapping on the plates immediately after illumination. Further checking after overnight recovery confirmed their continued viability. This suggests that exposure to this light level does not cause observable photo-toxic effects.

To further assess the impact of the excitation light on behavior, we measured on-food pumping in adult animals expressing the red fluorophore mCherry compared to the strain expressing YFP, which is used throughout the paper. If the impact of the wavelength is non-negligible, the red fluorophore mCherry (excitation centered at 587 nm) should result in fewer reversals or accelerations compared to YFP, as these responses are wave-length dependent [1,2,3]. As expected, we find an increase in the velocity for the green light exposed animals (Fig. S2.5A). However, we find that pumping rates between the two populations are not significantly different (Fig. S2.5B), suggesting that these light intensities do not affect pumping behavior. We note that the differential effects of excitation light on behavior should be taken into account when investigating the coupling between e.g., locomotory and feeding behaviors.
